# Quantitation of vector uptake reveals non-Poissonian transfection dynamics in *Plasmodium falciparum*

**DOI:** 10.1101/681981

**Authors:** Manuela Carrasquilla, Theo Sanderson, Ruddy Montandon, Julian Rayner, Alena Pance, Marcus Lee

## Abstract

The recurrent emergence of drug resistance in *Plasmodium falciparum* increases the urgency to genetically validate drug resistance mechanisms and identify new targets. Reverse genetics have facilitated genome-scale knockout screens in *Plasmodium berghei* and *Toxoplasma gondii*, in which pooled transfections of multiple vectors were critical to increasing scale and throughput. These approaches have not yet been implemented in *P. falciparum*, mainly because the extent to which pooled transfections can be performed in this species still remains unknown. Here we use next-generation sequencing to quantitate uptake of a pool of 94 barcoded vectors. The distribution of vectors in different transfections allowed us to estimate the number of barcodes and DNA molecules taken up by the parasite population. Dilution cloning showed that single transfected parasites routinely carry as many as seven episomal barcodes, revealing an intake of multiple vectors in a highly non-uniform fashion. Transfection of non-overlapping fluorescent proteins, which allowed us to follow the dynamics of the process, confirmed the tendency for parasites to take up multiple vectors from the early stages of transfection. This finding has important implications for how reverse genetics can be scaled in *P. falciparum*.

## Introduction

Reverse genetics is a key tool in the global effort to identify drug targets or resistance mechanisms, as well as to explore new biology. Technologies for genetic manipulation of organisms have advanced significantly in the last decade, particularly through site-specific nucleases such as Cas9 [1] that can be used to increase the efficiency of modification. However, using genetics to validate gene function in *Plasmodium falciparum*, the most virulent of the causative agents of human malaria, has been consistently challenging, for multiple reasons. The high AT content of its genome (>80% AT, up to ~90% in introns) makes generating large stable plasmids in *E. coli* difficult, and also limits the potential targets for Cas9 due to its requirement for an NGG protospacer adjacent motif (PAM) sequence, which are much rarer in *P. falciparum* genomic DNA than most eukaryotes. *P. falciparum* also has low transfection efficiencies compared with other *Plasmodium* species [2,3], despite attempts to generate more efficient protocols [4,5]. These constraints have limited progress in interrogating the genome of the parasite to uncover potential new drug targets and the roles of the many genes of unknown or poorly described function.

There are also specific challenges to the application of CRISPR/Cas9 technology, which has revolutionised genetic screening in many other eukaryotes [6]. Analysis of the *P. falciparum* genome [7] indicates that this organism lacks one of the two major DNA repair mechanisms, non-homologous end joining (NHEJ). In principle, this should provide an advantage for genome editing, as the introduction of a double strand break in the parasite DNA coupled with homology templates that provide the desired modification should result in consistent homology-directed repair without competing error-prone events. As a result, the application of CRISPR/Cas9 in *P. falciparum* has relied on donor templates and has been used to knockout genes [8,9], and validate key drug resistance mechanisms [10–12]. However the absence of NHEJ, coupled with the low transfection efficiency noted above, has made systematic large-scale gRNA-based gene disruption screens that have been revolutionary in other organisms elusive so far [6,13]. Newly developed CRISPRi and CRISPRa approaches [14–16], which use a nuclease-dead version of Cas9 as a DNA-binding protein to either repress or activate gene expression, might allow multiplexed screening even in the absence of NHEJ machinery. However, such approaches will rely on achieving relatively high transfection efficiencies, and an understanding of how many unique plasmids (containing different gRNAs) are taken up by each parasite.

While *P. falciparum* co-transfection is routinely performed with two or three plasmids that each have unique selectable markers, to our knowledge the dynamics of *P. falciparum* transfection has not been studied with techniques that allow the multiplicity of vector uptake to be accurately quantitated. We sought to measure the propensity of *P. falciparum* to take up multiple plasmids by transfecting pools of vectors that could be distinguished either on the basis of sequence or on the fluorescent properties of their encoded products. We then examined both the proportion of these vectors taken up by parasites in bulk culture, and how these events were distributed within cloned parasite populations. In one set of experiments we co-transfected 94 vectors that each encoded a unique DNA barcode, and used next generation sequencing to quantitate their uptake in the bulk parasite population as well as in cloned transfected parasites. In a second approach, we transfected a combination of expression vectors encoding fluorescent proteins, and followed their expression in the transfected parasite population over time. The data from both these approaches help us to understand the complexity and dynamics of transfection, and have important implications for the design of future *P. falciparum* genetic experiments.

## Results

### Transfection of a pooled barcode library

In order to examine the diversity of vector uptake after transfection, we created a large pool of plasmids that differed only in a short 11 bp barcode, allowing identification of each plasmid uniquely by next generation sequencing, but minimising differences in sequence that may bias their propagation in the parasite. The pool of vectors was created by first amplifying 96 unique DNA barcodes within a short 120 bp cassette [17] and assembling them, in a single Gibson reaction, into a vector backbone that contained the h*DHFR* selectable marker, allowing for positive selection (Fig. 1a). This vector pool was transfected into standard *P. falciparum* laboratory strains 3D7 and Dd2, and constant drug pressure with WR99210 was applied from one day post-transfection. Once parasites became visible by Giemsa stain, the cultures were expanded, and parasite genomic DNA was extracted. Barcodes were amplified by PCR and quantitated by Illumina sequencing (Fig 1b-c) using the barcode sequencing (Bar-Seq) method pioneered in *P. berghei* [18]. The starting vector pool was also sequenced to allow for comparison of barcode distributions before and after transfection. The transfection was repeated on five independent occasions.

**Figure 1.**
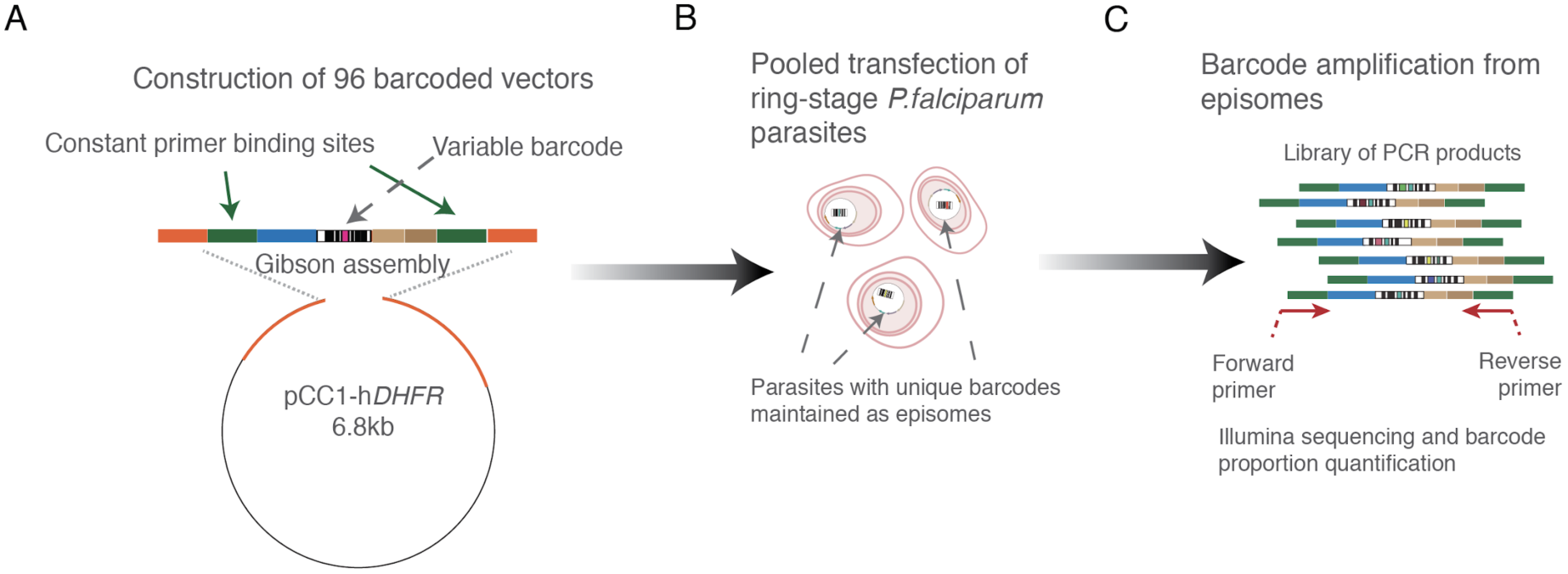
Graphical representation of the plasmid assembly method using an amplicon containing 96 unique barcodes amplified from a library of pGEM Knockout vectors [2]. **A.** Amplicon assembly of all 96 barcodes into the pCC1-hDHFR transfection vector was performed in a single Gibson reaction, and the bacterial transformation was used to directly seed a large overnight culture for midiprep plasmid preparation. This starting pool was examined by Bar-Seq to confirm representation of barcodes (see Fig. 2). **B.** Pools of 94 constructs were used as DNA input for *P. falciparum* ring-stage transfection. **C.** Parasite genomic DNA was extracted from both the bulk culture and individual clones for amplification of the barcode from episomes for Next Generation Sequencing and barcode quantification.

Ninety four of the 96 barcodes were represented in the input pool used for transfection (Fig. 2, Input), albeit at differing frequencies. However, examination of the bulk population of transfected parasites indicated that not all barcodes had been taken up by parasites and maintained as episomes. To quantify the number of unique barcodes taken up, as well as to estimate the total number of plasmids acquired, for each barcode we calculated the difference between its log-ratio in the input and in the final population of parasites. This resulted in a bimodal distribution as shown (Supplementary Material 1), which we took to represent a superposition of two normal distributions: one representing barcodes successfully taken up by parasites and another representing barcodes that were not. In each case we fitted a mixture of these two distributions to estimate the number of unique barcodes in each transfection. By simulating random sampling of barcodes with the relative proportions found in the input, we were able to estimate the number of molecules taken up in each transfection, which ranged from 9 to 130 (Supplementary Material 1). This translated to a range of 9-66 unique barcodes represented in the bulk populations (Table 1).

**Table 1.** Modelled values for the number of barcodes taken up in each transfection, and the expected number of molecules of DNA acquired that this corresponds to, given the possibility that one barcode could be taken up multiple times.

| Transfection | Unique barcodes | Molecules of DNA taken up |
| --- | --- | --- |
| 1 | 44 | 64 |
| 2 | 66 | 130 |
| 3 | 25 | 31 |
| 4 | 19 | 22 |
| 5 | 9 | 9 |

**Figure 2.**
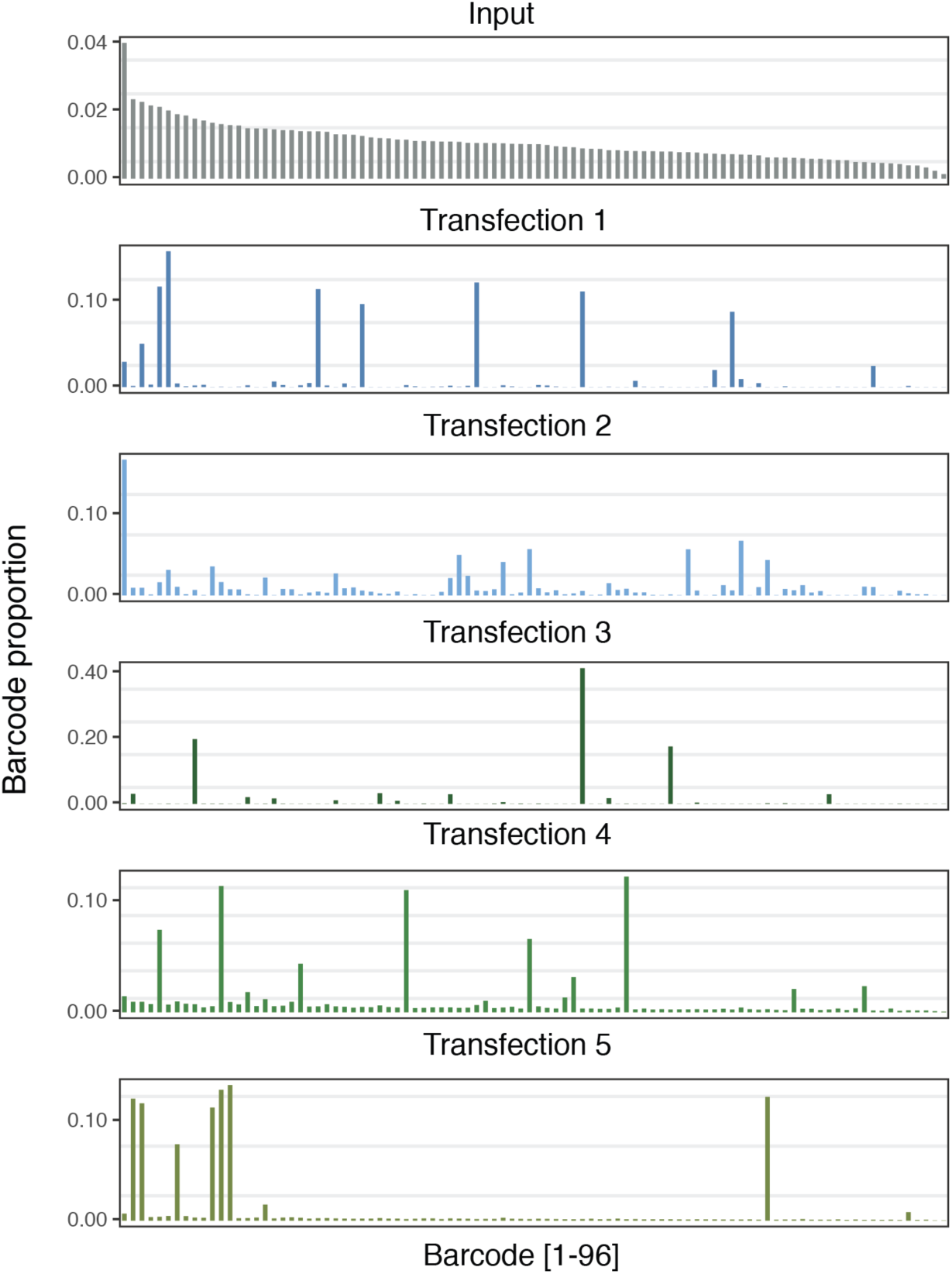
Transfection of a pool of 94 uniquely barcoded vectors. Individual barcodes are represented on the x-axis, and the data is sorted by the most abundant barcodes present in the input pool (first panel, grey). The y-axis represents the proportion of each barcode in the total pool, per individual transfection as obtained with Next Generation Sequencing.

### Co-transfection results in uptake of multiple vectors by a small proportion of parasites

The fact that 9-66 vectors, from an available pool of 94, could be recovered from each transfection might suggest a potential for multiplexed screening. However, *P. falciparum* parasites are known to be able to take up at least two plasmids when they are transfected together, as evidenced by the fact that co-transfection with two different selectable markers has been used to co-localise differentially tagged proteins [19,20], and the application of CRISPR/Cas9 to date has routinely relied on transfection with one plasmid containing the Cas9 endonuclease and another containing the homology repair construct [8]. While the efficiency of CRISPR/Cas9 editing in *P. falciparum* is notoriously variable, it does confirm that *P. falciparum* parasites do repeatedly take up more than one plasmid at the same time. To assess the distribution of barcodes in the parasite population and establish the extent to which individual parasites take up multiple plasmids, we cloned single parasites by limiting dilution from the bulk culture recovered from transfection 5 (Fig. 3a). Barcode sequencing of ten clones showed that all contained multiple barcodes, with 4 to 7 barcodes detected per clone (Fig. 3b). Furthermore, only two different combinations of barcodes were observed from the clones (clone types 1 and 5), with most of the clones having the same barcodes represented, although at different relative levels. These observations reveal that even though transfection efficiency of *P. falciparum* is very low (Fig. 3a), there is clearly a non-random distribution of plasmid uptake, and a strong preference for individual parasites to take up multiple plasmids.

**Figure 3.**
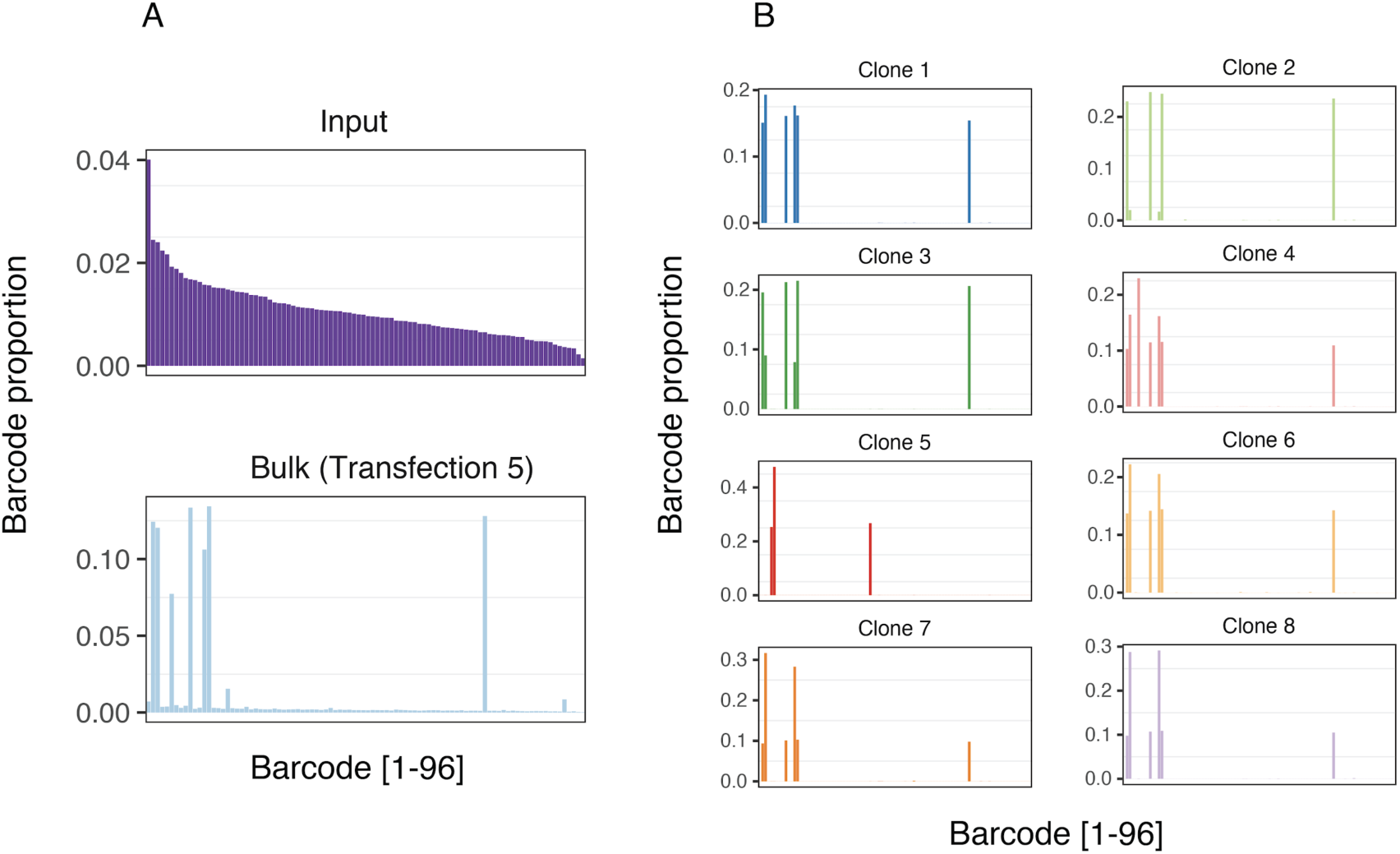
Measurement of the complexity of episomes in individual parasite clones obtained with limiting dilution from the least complex transfection (5). **A.** The top panel shows the input plasmid pool and the bottom panel shows the complexity in the bulk population. **B.** Barcode sequencing of eight clones revealed two clone-types were obtained, with all but clone 5 representing a single clone type. The y-axis is as described for Fig. 2.

### Dynamics of transfection with fluorescent proteins confirms uptake of multiple plasmids from the early stages of transfection

To further investigate these results, as well as generate a resource of fluorescent parasites for future parasite competition assays, we constructed a set of vectors for episomal expression of a range of fluorescent proteins. A variety of genes encoding proteins with diverse excitation/emission characteristics (Table 2) were cloned into expression vectors under the control of the *P. falciparum* calmodulin promoter, and with a blasticidin selection cassette in the vector backbone. The compatible fluorochromes tagBFP, MiCy (Midori-ishi Cyan) and mCherry were chosen to transfect NF54 parasites (Fig. S1 in the Supplementary Information). The use of fluorescent proteins allowed the kinetics of the transfected populations to be followed by flow cytometry (Fig. 4a and Fig. S2 in the Supplementary Information). Vectors were taken up in every possible combination by the population of parasites, and maintained throughout the entire time course of observation. From the earliest time points, we observed that simultaneous expression of multiple fluorochromes was a commonplace occurrence. These data confirm the non-independence of plasmid uptake observed using the barseq-based methodology, and the appearance of such parasites so soon after transfection suggests a phenomenon occurring early during transfection. Furthermore, the use of fluorescent markers enabled qualitative exploration by microscopy, where parasites carrying one, two, and all three vectors were all visible (Fig 4b-e). These direct observations allowed us to rule out technical explanations of our results, such as the possibility of compromised dilution cloning in the barseq experiment, and point to the uptake of multiple plasmid molecules per cell as the predominant transfection event.

**Table 2:** Excitation/emission values for fluorochromes used.

| Fluorochrome | Excitation (nm) | Emission (nm) | Laser |
| --- | --- | --- | --- |
| tagBFP | 399 | 456 | 405 |
| Midori-ishi Cyan | 472 | 495 | 488 |
| mCherry | 587 | 610 | 561 |

**Figure 4.**
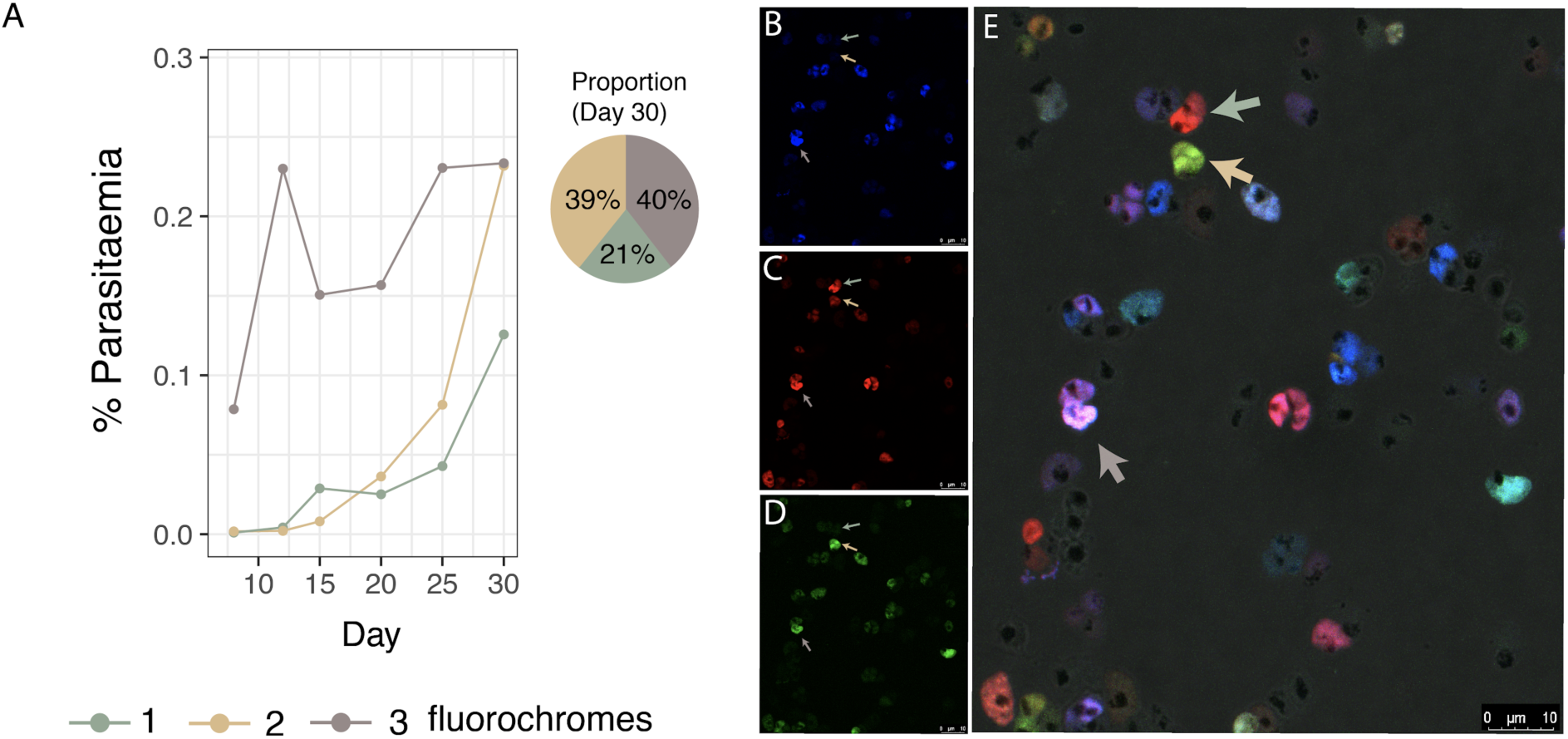
**Time course of transfected parasites expressing different combinations of fluorochromes, confirming multiple plasmid uptake, A.** The proportion of parasites with one (green), two (yellow) or all three (grey) fluorochromes followed over 30 days, following transfection with MiCy, mCherry and BFP. Blasticidin selection (2µg/mL) was applied the next day after transfection for one week. A representative experiment is shown, with two independent replicates shown in Fig. S2. **B-E.** Microscopy of tagBFP (B), mCherry (C) and Midori-ishi Cyan (D) fluorochromes transfected together (merge including brightfield shown in panel E). Green, yellow and grey arrows label exemplar parasites simultaneously expressing one, two or three fluorochromes, respectively.

## Discussion

Experimental genetics has had a long track record of revealing new biology in *P. falciparum*, and recent developments have yielded multiple new promising tools to even more exquisitely manipulate the genome of this globally significant parasite [21]. However, all the technologies available depend on the capacity to introduce exogenous DNA into the parasite, and this remains a significant limitation for the field. Two decades after the first transfections it is still unclear what factors limit the efficiency with which DNA can be delivered to *P. falciparum* through transfection, and why this efficiency appears to be much lower than in other *Plasmodium* species [3,22]. Our data with the complex pool of barcoded plasmids allowed us to calculate the number of DNA molecules delivered per transfection. We estimate that ~6 × 10^7^ parasites were placed in the transfection cuvette, and collectively took up between 9 and 130 molecules of DNA upon electroporation for a maximum transfection efficiency of ~1 × 10^−6^, assuming independent plasmid uptake modelled under a Poisson distribution. This underlines the very low transfection efficiency of *P. falciparum* - by contrast with *Plasmodium knowlesi* or *Toxoplasma gondii* [3,23]. Our data also yields another unexpected hurdle to large-scale screening in *P. falciparum*. If each of these 9-130 uptake events were independent, one would expect each of these molecules to be found in a different parasite. Under this model of independent uptake events, the chances of a single parasite taking up 2 molecules would be ~1 × 10^−12^; and given that there are only ~5×10^7^ fewer parasites in the cuvette, we would never expect to see such a cell. In fact, our data from clonal dilution and fluorescence experiments suggest that the vast majority of transfection events involve the uptake of multiple molecules, and that parasites routinely acquire as many as seven. The appearance of transfected parasites expressing more than one fluorescent protein from the very earliest stages of detection suggests that this phenomenon is driven by the uptake of multiple plasmids during transfection, rather than the exchange of plasmids between parasites post-transfection, for example through microvesicles [24,25], although it should be acknowledged that the dynamics or frequency of such exchange is not well established. Regardless of the mechanism, the implications for the design of *P. falciparum* pooled transfection experiments is clear - that parasites harbouring multiple plasmids are the norm, rather than the exception.

These data concur with the success that *Plasmodium* researchers have had with two plasmid transfection approaches, for instance in co-localisation experiments [26] or for the delivery of Cas9 and donor plasmids [8,9], and suggest that these should not be significantly less efficient than one plasmid approaches. These results suggest that the actual transfection efficiency for *P. falciparum* is even less than ~1 × 10^−6^. They also raise caveats about the approach of transient transfection of a luciferase reporter, a common method used to optimise transfection conditions [27]. A luciferase assay in essence measures the number of plasmids taken up by a culture, but has little power to assess how these plasmids are distributed throughout the parasite population, and so current techniques may not have been optimised for the number of parasites taking up exogenous DNA.

There are a number of possible hypothetical bottlenecks in *Plasmodium* transfection. Our data suggest that availability of DNA is not likely to be a crucial limitation, and it is possible that lower concentrations of DNA might even achieve superior results. It also seems likely that the use of “two plasmid” systems, for instance for supplying a donor and a Cas9 vector, are unlikely to lead to significantly lower efficiencies than one plasmid systems with current transfection conditions. A previous similar analysis carried out in Jurkat cells using fluorescent markers provided a result much closer to a Poissonian process, with cells differing in their susceptibility to transfection by at most a factor of three [28]. One potential explanation for our data is that only a few *P. falciparum* cells within a culture are actually competent to take up plasmids, and efforts to identify the nature of these rare parasites, and also what distinguishes *P. falciparum* from other *Plasmodium* species with much higher transfection efficiencies may yield insights [3,22]. Further understanding these dynamics and the factors that control them will be particularly important for the development of multiplexed CRISPR-interference or CRISPR-activation screens, where low multiplicities of transfection will be crucial. This knowledge will be fundamental to the exploration of the parasite’s genome and the unravelling of gene function as well as host-parasite interactions.

## Methods

### Generating fluorochrome-expressing vectors

Midori-ishi Cyan (acquired from Addgene), tagBFP and mCherry sequences were amplified by PCR using specific primers with restriction sites for AvrII for the forward and XhoI for the reverse primers. The fragments were ligated into the pDC2 *Plasmodium* expression vector [29] containing a PfCAM promoter and a Blasticidin-S deaminase selectable marker, for drug selection at 2 µg/ml Blasticidin.

### Generation of library of barcoded plasmids

The templates for barcode amplification were derived from PlasmoGEM resource pGEM vectors (https://plasmogem.sanger.ac.uk), provided as glycerol stocks. 96 different bacterial stocks each possessing an individual barcode vector were incubated individually in deep 96-well plates (maximum volume 2 mL) overnight at 30 °C, in TB medium with 30 µg/mL of kanamycin. After each bacterial culture reached an appropriate optical density, and assuming similar growth rates between them, the cultures were pooled and the library of 96 vectors was extracted using a Macherey Nagel midi prep kit. Amplicons contained in the 96 different vectors were amplified directly from the pool using primers p212 (CAATTAATGATGTATACCGCCTTCAATTTCGATGGGTAC) and p219 (CTAAGAAGGTTATAGAGGCGTAATTCGTGCGCGTCAG), which were designed to amplify a 120 bp barcode amplicon with sequence overlap to a NheI/NcoI digested *P. falciparum* vector pCC1 to permit Gibson assembly (Fig. 1a). The final assembled vector containing all possible 96 barcodes cloned into pCC1 was transformed into One-Shot TOP10 competent *E.coli* and plated onto LB plates with ampicillin. Colonies were pooled into a large culture and a midiprep was performed to generate the transfection pool.

### Parasite cultures

Parasites were propagated at 37 °C and with a gas mix of 1% O2, 3% CO2 and 96% N. All parasite strains were routinely cultured in O+ Red Blood Cells (RBCs) provided by anonymous healthy donors from the National Health Services (NHS), in standard parasite culture media essentially as described in [30]. Synchronization of cultures was performed using 5% sorbitol in water as in [31].

### Transfection of parasites

For pooled barcode transfections, plasmid DNA (50 µg per pool) was resuspended in 100 µl of buffer P3 (Lonza) with 4 µl ATP (625 mM). Cultures containing high parasitemia (approximately 10%) at mostly ring stage were centrifuged and 100 µl of packed RBCs per transfection were washed with cold cytomix, resuspended in P3 buffer with DNA and ATP and electroporated with a Lonza Nucleofector 4D using the programme P3/CM-150. The fluorochrome vectors were transfected following the same procedure using 20 µg of each construct and a culture of NF54 parasites.

### Library preparation for Next Generation Sequencing

Genomic DNA was extracted using a Qiagen Blood and Tissue Kit after parasites reached 5% parasitaemia. Once gDNA was extracted and measured, a nested PCR reaction targeting the constant flanking regions of the barcode was performed. 50ng of gDNA was used for amplification using p1356 (TCGGCATTCCTGCTGAACCGCTCTTCCGATCTGTAATTCGTGCGCGTCAG) and p1357 (ACACTCTTTCCCTACACGACGCTCTTCCGATCTCCTTCAATTTCGATGGGTAC) containing adapters for Illumina. Paired-end index primers (Illumina Nextera) followed on a secondary PCR. All reactions were performed using a 2X KAPA Hot-Start Master Mix (Kapa Biosciences). A PCR purification step was performed only on the second PCR using a Macherey Nagel PCR purification kit and eluted DNA was measured, multiplexed and diluted to a final concentration of 4 nM. Samples were loaded onto an Illumina MiSeq sequencer, using a MiSeq Reagent Kit v2 (300 cycle). They were loaded at a low cluster density (< 400 k cluster density), and 50% of PhiX was spiked in, as described in Gomes et al. (2015) for low complexity libraries [18].

### Barcode counting and analysis

Barcode counting was performed as in [18]. Briefly, raw reads coming out of the Illumina MiSeq, represented by unique index tags, were separated and analysed with a script that identified correct flanking sequences and counted exact matches of unique barcodes between these constant regions. Barcode counts for the experiments shown in Figures 2 and 3 are provided in the Supplementary Information.

### Cloning of parasites by limiting dilution

To clone transfected parasites, we used limiting dilution cloning using a modified version of [32]. Parasites were diluted into a 96-well plate at 0.5-0.8 parasites per well at 1.8% haematocrit, expecting approximately 50% of the plate to contain clonal parasites. Detection of positive wells was first performed using the DNA stain SYBR Green, discarding either empty wells or those with much higher fluorescence than the average, indicating possibly more than one parasite was inoculated at seeding.

### Flow Cytometry

After transfection, the parasite cultures were followed every 4 to 5 days, when samples were taken, fixed in 4% paraformaldehyde for 20 minutes, washed and resuspended in PBS. Excitation/emission values for the fluorochromes used is listed in Table 2. Gating was established with blood alone and untransfected parasite cultures to measure fluorescence with a BD LSRFortessa cell analyser (BD Biosciences, Oxford UK) collecting 200,000 events per sample. FlowJo (7.6.5) was used for the analysis and the gating strategy, after removing doublets, is shown in Fig. S3 in the Supplementary Information.

### Microscopy

Standard blood smear microscopy was performed to determine parasitemia. In brief, a small aliquot of culture was smeared on a glass slide, fixed with 100% methanol and stained with a 10% Giemsa solution (Sigma-Aldrich). Fluorescent parasites were imaged on a Leica TCS SP8 confocal microscope (Leica Microsystems). Schizonts of each transfected culture were purified on a 63% percoll gradient and fixed with fresh 4% paraformaldehyde for 20 minutes. The schizonts were placed on slides, dried and mounted with ProLong Diamond antifade mountant (Thermo Fisher) overnight at 4°C.

### Data availability

All data necessary to perform the analysis of the BarSeq transfections are available (Supplementary Material 1), as well as the data on the barcode complexity of the bulk culture and isolated clones (Fig. 2, Fig. 3, and Supplementary Material 2).

## Supporting information

Supplemental Information all

## Acknowledgements

We would like to thank members of the Lee, Rayner, and Billker labs for productive discussions. We are grateful to the staff in Sanger Scientific Operations for their support with sequencing. This work was supported by funding from Wellcome (206194).

## Authors contributions

M.C., A.P., J.R. and M.L. conceived the experiments. M.C. performed the barcode cloning and transfection, and with T.S. analysed the sequencing data. A.P. performed the fluorescent protein cloning, transfection and flow cytometry, with assistance from R.M. All authors contributed to writing the paper.

