## Supplemental Information all for "Quantitation of vector uptake reveals non-Poissonian transfection dynamics in *Plasmodium falciparum*"

2 Present address: Department of Immunology and Infectious Diseases, Harvard T.H. Chan School of Public Health, Boston, U.S.A.

3 Present address: Wellcome Centre for Human Genetics, Oxford, UK.

4 Present address: Cambridge Institute for Medical Research, Cambridge, UK.

### Supplementary Figure 1

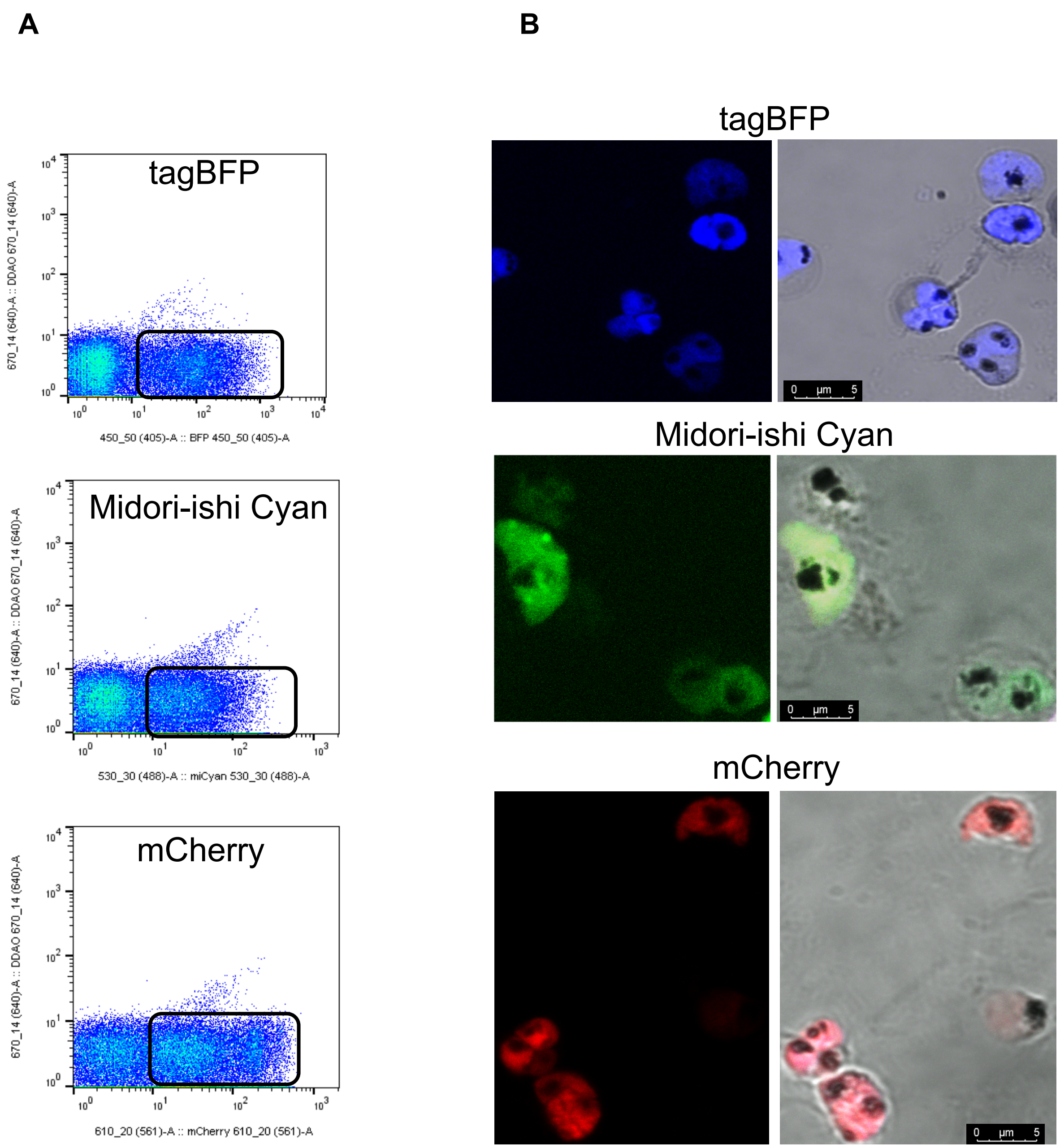


**Figure S1.** *Plasmodium falciparum* NF54 parasites were transfected with individual fluorochromes tagBFP (blue), Midori-ishi Cyan (green) or mCherry (red) (ex/em listed in Table 2), which were expressed from episomal plasmids under blasticidin selection (2 µg/mL). **A.** Recovered parasites after transfection were visualised by flow cytometry and to determine parasitemia, black square shows the fluorescent population for the different proteins. **B.** Fluorescent cells were confirmed with confocal microscopy.

### Supplementary Figure 2

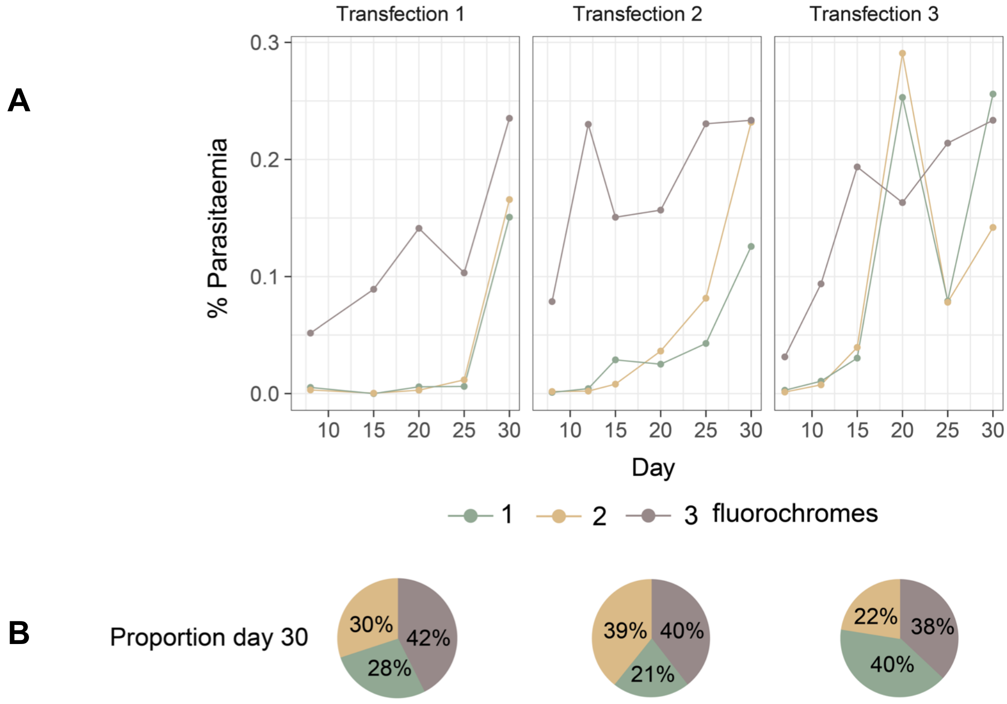


**Figure S2. A.** Dynamics of three independent transfections over the course of 30 days. Parasites were co-transfected with the three expression plasmids for the tagBFP, Midori-ishi Cyan or mCherry fluorochromes, and were followed from day 10 post-transfection with flow cytometry. **B.** Proportion of expressing one, two or three fluorochromes on day 30, measured by flow cytometry

### Supplementary Figure 3

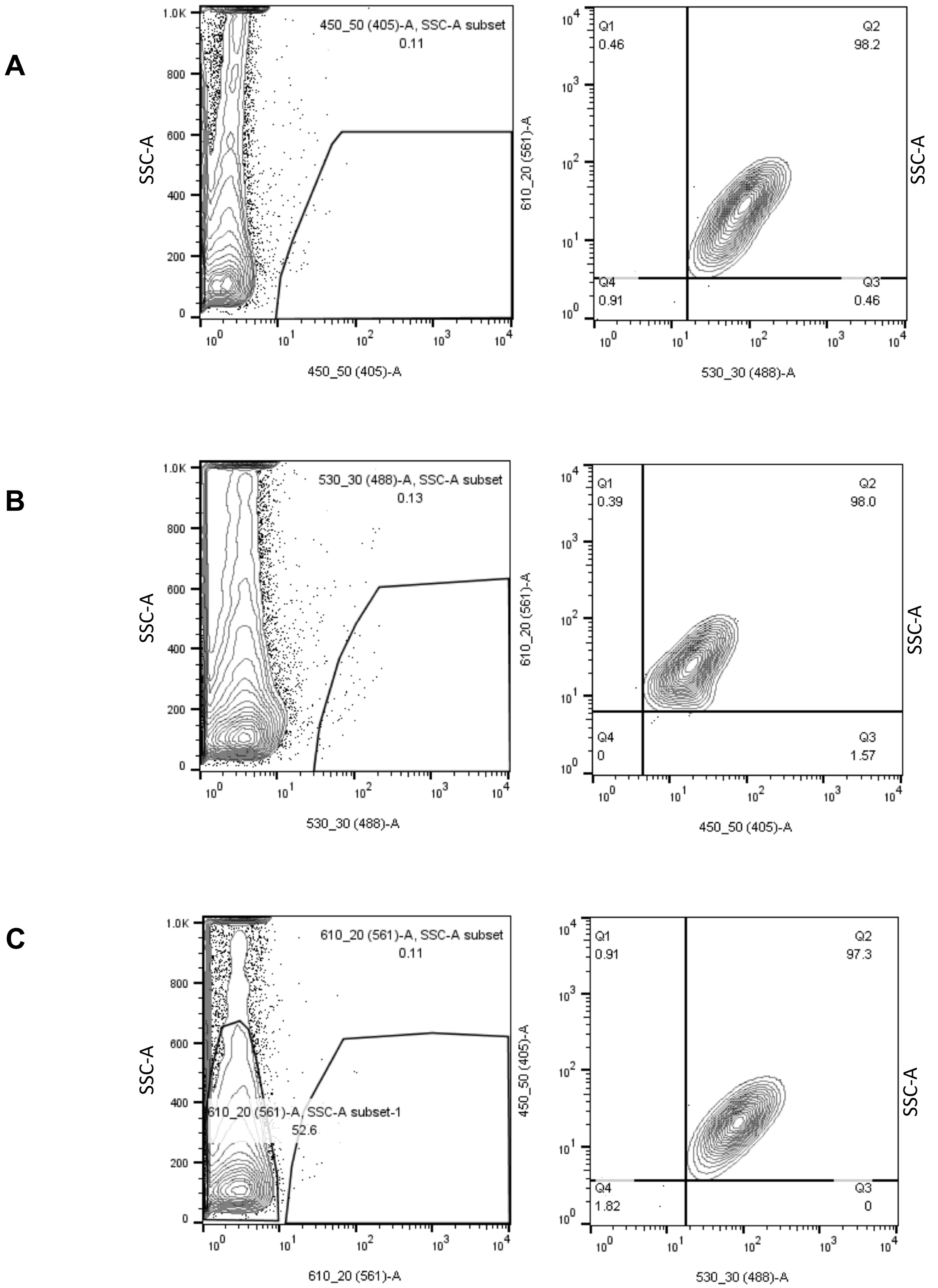


**Figure S3.** Gating strategy for the flow cytometry assessment of fluorescent parasites. Once the population is determined and doublets are removed the fluorescent events in the culture are quantified using the appropriate lasers and filters (Table 2, Methods). Each fluorochrome is quantified and the fluorescent population is then analysed under the excitation/emission conditions of the other two. The resulting populations for each combination are averaged.

### Analysis of episomal barcodes distributions

We are first going to fit distributions to estimate the number of unique barcodes in each transfection. First we load the data:

```
library(knitr)
library(kableExtra)
library(tidyverse)
library(mixtools)
dat<-read_csv("Supp_transall.csv")
narrow<- dat %>% gather("type","val",-barcode)
```

Now we convert everything to log2 format, and work out for each barcode the difference between its value in the input and its value in each of the transfections. We plot the distribution of these differences.

```
narrowlog <- mutate(narrow,val=log2(val+0.5))
```

```
## Warning: package 'bindrcpp' was built under R version 3.4.4
```

```
input<-filter(narrowlog,type=="input")
noninput<-filter(narrowlog,type!="input")
both<-inner_join(noninput,input,by="barcode") %>% mutate(diff=val.x-val.y)
ggplot(both,aes(x=diff,color=type.x))+geom_density() +labs(x="Difference (in log-base-2)",y="Density", 
```

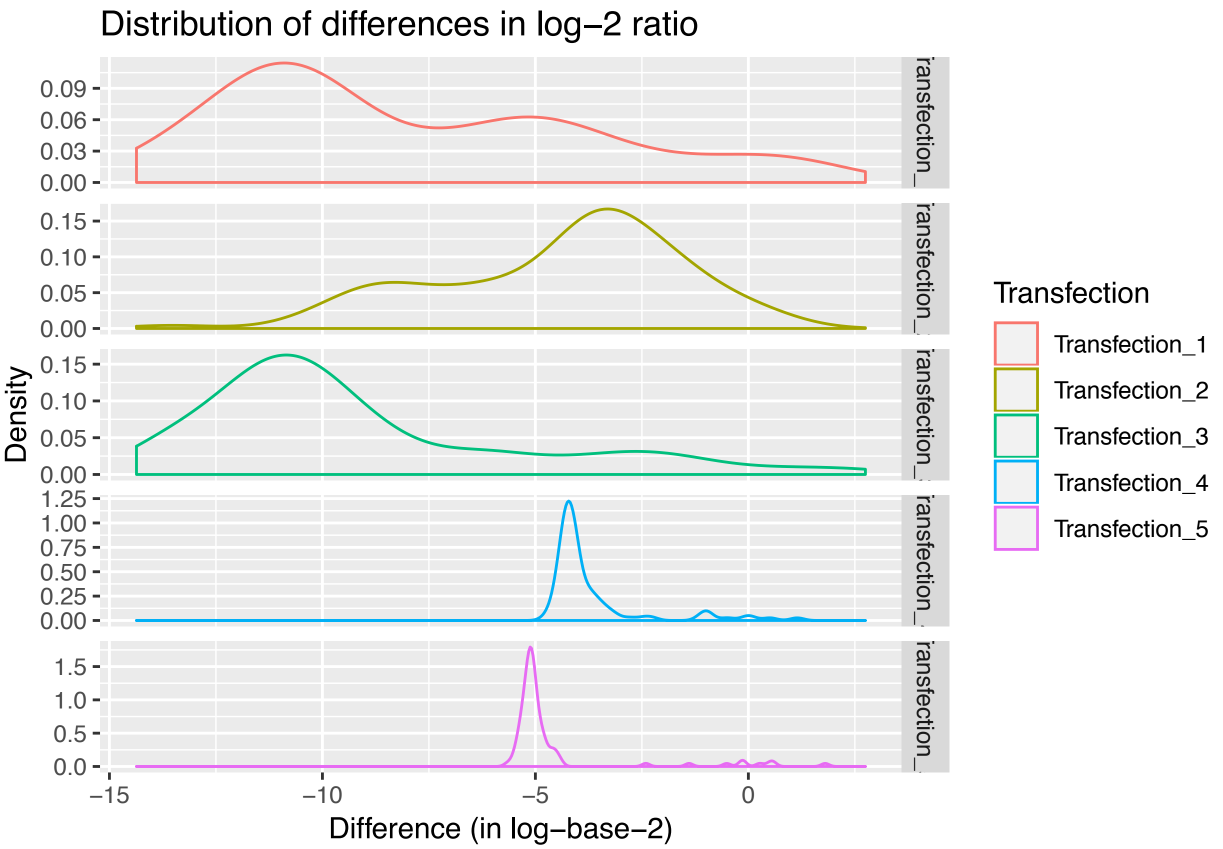


Since all barcodes should have equal fitness, variance in this plot should arguably only be explained by two factors: stochasticity in whether a barcode ever made it into the population of parasites, and then stochasticity in the growth of those parasites. There appears to be a bimodal distribution.

We model this as the sum of two normal distributions, in the expectation that the more distribution with the low mean represents complete failure to establish a transfectant and the more positive distribution represents establishment of a transfectant. We deconvolute this mixture below.

```
dnormWithLambda<-function(x,mean,sd,lambda){
  results<-dnorm(x,mean,sd)
  return(results*lambda)
}
splitUp <- function(data){
  mod<-normalmixEM(data$diff,epsilon=1e-20,maxrestarts=500)

  lambdas=mod$lambda
  mus=mod$mu
  sigmas=mod$sigma

  if (mus[1]>mus[2])
  {
    lambdas=rev(lambdas)
    mus=rev(mus)
    sigmas=rev(sigmas)

  }

  p<-ggplot(data,aes(x=diff))+geom_density(color="red")+stat_function(color="blue",linetype=2,fun = dno
 print(p)
        return(data.frame(lambda=lambdas[2]))

}
library(tidyverse)
resultsFirst<-both %>% group_by(type.x) %>% do(splitUp(.)) %>% dplyr::mutate(numberOfBarcodes=round(94*
```

```
## number of iterations= 35
```

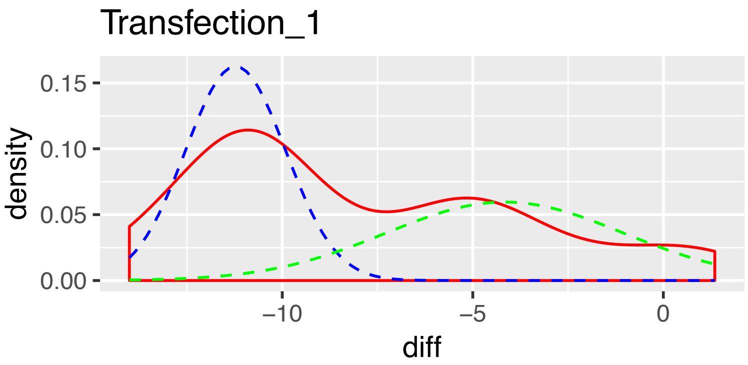


```
## number of iterations= 150
```

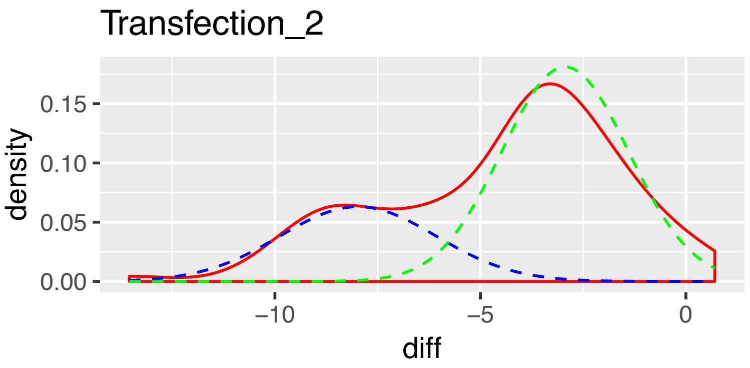


```
## number of iterations= 78
```

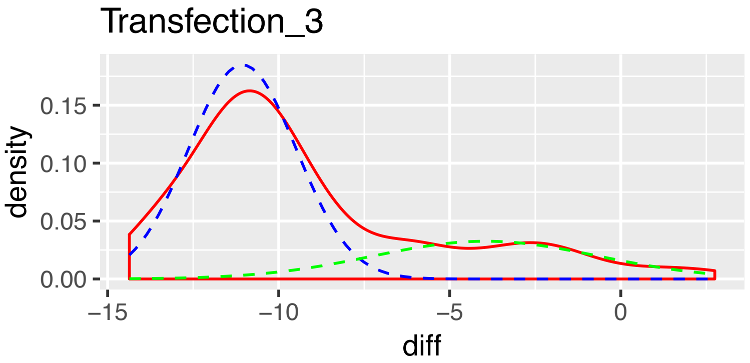


```
## number of iterations= 74
```

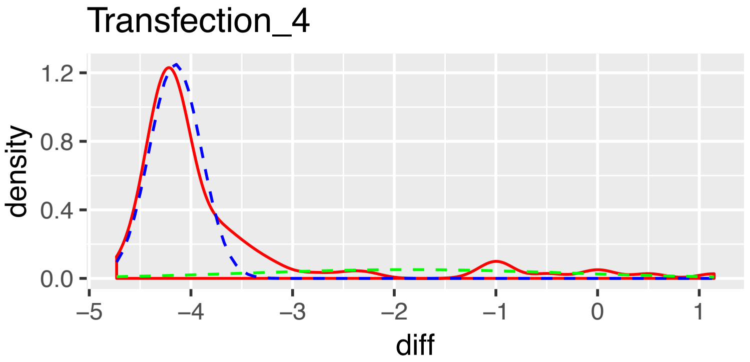


```
## number of iterations= 11
```

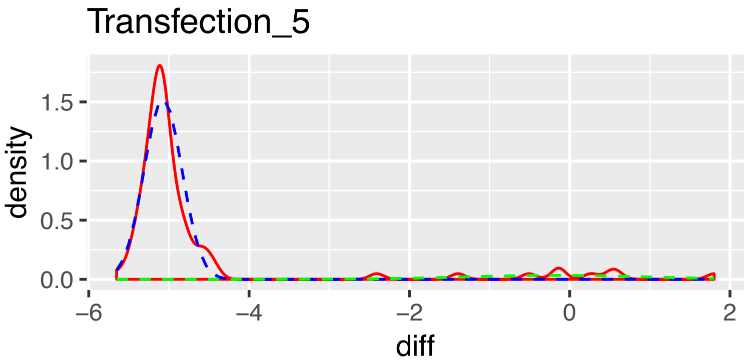


```
resultsFirstDisp <- resultsFirst
resultsFirst
```

```
## # A tibble: 5 x 3
## # Groups:   type.x [5]
##   type.x          lambda numberOfBarcodes
##   <chr>            <dbl>            <dbl>
## 1 Transfection_1 0.464               44
## 2 Transfection_2 0.699               66
## 3 Transfection_3 0.269               25
## 4 Transfection_4 0.204               19
## 5 Transfection_5 0.0958               9
```

```
colnames(resultsFirstDisp)=c("Transfection","Lambda","Unique Barcodes")
kable(resultsFirstDisp,format="latex",booktabs=T)
```

| Transfection | Lambda | Unique Barcodes |
|---|---|---|
| Transfection_1 | 0.4641848 | 44 |
| Transfection_2 | 0.6994108 | 66 |
| Transfection_3 | 0.2691667 | 25 |
| Transfection_4 | 0.2038329 | 19 |
| Transfection_5 | 0.0957712 | 9 |

We now have an estimate of the number of unique barcodes in each transfection.

We can now ask another question. Given the proportion of various barcodes in the input, if we simulate a certain number of barcodes being taken up by parasites, then how many distinct barcodes would we expect to recover (given that one batcode might be taken up more than once)?

```
input2<- narrow %>% filter(type=="input") %>% mutate(prop=val/sum(val))
simulate <- function(numberTakenUp){

answer=median(replicate(100,length(unique(sample(x = 1:nrow(input), numberTakenUp, replace = T, prob = 
return(answer)
}
numberTakenUp<-1:500
numberOfBarcodesRecovered<-sapply(numberTakenUp,simulate)
df<-data.frame(numberTakenUp=numberTakenUp,numberOfBarcodesRecovered=numberOfBarcodesRecovered)
ggplot(df,aes(x=numberTakenUp,y=numberOfBarcodesRecovered))+geom_line()+labs(x="Number of molecules tak
```

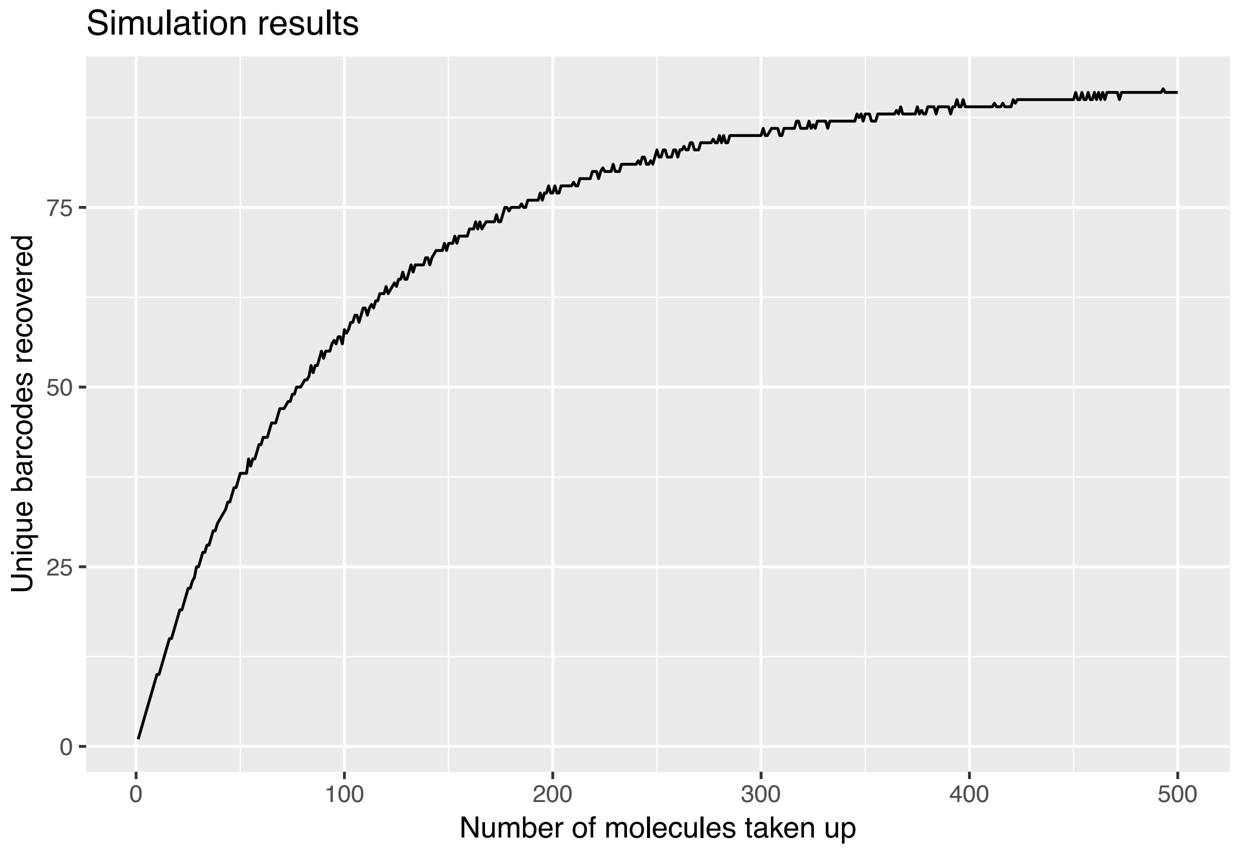


```
df <- df %>% group_by(numberOfBarcodesRecovered) %>% summarise(numberTakenUp=mean(numberTakenUp))
```

We can now use this data to estimate how many molecules of DNA were taken up in each transfection.

```
combination<-inner_join(resultsFirst,df,by=c("numberOfBarcodes"="numberOfBarcodesRecovered"))
colnames(combination)=c("Transfection","Lambda","Unique Barcodes","Molecules of DNA taken up")
kable(combination,format="latex",booktabs=T)
```

| Transfection | Lambda | Unique Barcodes | Molecules of DNA taken up |
|---|---|---|---|
| Transfection_1 | 0.4641848 | 44 | 64.0000 |
| Transfection_2 | 0.6994108 | 66 | 130.6667 |
| Transfection_3 | 0.2691667 | 25 | 29.5000 |
| Transfection_4 | 0.2038329 | 19 | 21.5000 |
| Transfection_5 | 0.0957712 | 9 | 9.0000 |

We conclude that in these transfections parasites took up and stabilised 9-130 DNA molecules, depending on the transfection.
